# Complex Structural Variants Resolved by Short-Read and Long-Read Whole Genome Sequencing in Mendelian Disorders

**DOI:** 10.1101/281683

**Authors:** Alba Sanchis-Juan, Jonathan Stephens, Courtney E French, Nicholas Gleadall, Karyn Mégy, Christopher Penkett, Kathleen Stirrups, Isabelle Delon, Eleanor Dewhurst, Helen Dolling, Marie Erwood, Detelina Grozeva, Gavin Arno, Andrew R Webster, Trevor Cole, Topun Austin, Ricardo Garcia Branco, NIHR BioResource, Willem H Ouwehand, F Lucy Raymond, Keren J Carss

## Abstract

Complex structural variants (cxSVs) are genomic rearrangements comprising multiple structural variants, typically involving three or more breakpoint junctions. They contribute to human genomic variation and can cause Mendelian disease, however they are not typically considered during genetic testing. Here, we investigate the role of cxSVs in Mendelian disease using short-read whole genome sequencing (WGS) data from 1,324 individuals with neurodevelopmental or retinal disorders from the NIHR BioResource project. We present four cases of individuals with a cxSV affecting Mendelian disease-associated genes. Three of the cxSVs are pathogenic: a *de novo* duplication-inversion-inversion-deletion affecting *ARID1B* in an individual with Coffin-Siris syndrome, a deletion-inversion-duplication affecting *HNRNPU* in an individual with intellectual disability and seizures, and a homozygous deletion-inversion-deletion affecting *CEP78* in an individual with cone-rod dystrophy. Additionally, we identified a *de novo* duplication-inversion-duplication overlapping *CDKL5* in an individual with neonatal hypoxic-ischaemic encephalopathy. Long-read sequencing technology used to resolve the breakpoints demonstrated the presence of both a disrupted and an intact copy of *CDKL5* on the same allele; therefore, it was classified as a variant of uncertain significance. Analysis of sequence flanking all breakpoint junctions in all the cxSVs revealed both microhomology and longer repetitive sequences, suggesting both replication and homology based processes. Accurate resolution of cxSVs is essential for clinical interpretation, and here we demonstrate that long-read WGS is a powerful technology by which to achieve this. Our results show cxSVs are an important although rare cause of Mendelian disease, and we therefore recommend their consideration during research and clinical investigations.

## Main

Structural variants (SVs) are a major source of variation in the human genome and collectively account for more differences between individuals than single nucleotide variants (SNVs).^1–3^ The canonical forms include balanced and unbalanced SVs, such as inversions, insertions, translocations, deletions and duplications. More complex rearrangements are typically composed of three or more breakpoint junctions and cannot be characterised as a single canonical SV type. These are known as non-canonical or complex SVs (cxSVs).^4; 5^

Several previous studies have reported clinically relevant cxSVs in individuals with Mendelian disorders. For example, a duplication-triplication-inversion-duplication was found at the *MECP2* and *PLP1* loci in individuals with Rett syndrome,^6^ and a duplication-inversion-terminal deletion of chromosome 13 was present in foetuses with 13q deletion syndrome,^7^ among others.^8–10^ Recently, pathogenic cxSVs associated with autism spectrum disorder (ASD) and neuropsychiatric disorders have also been reported.^11; 12^ Whole-genome sequencing (WGS) studies have shown that cxSVs are considerably more abundant and diverse than had been previously appreciated, representing an estimated 2% of the SVs in the human genome, and each human genome contains on average 14 cxSVs.^11^ The presence of multiple types of cxSVs has also been independently observed in several other studies.^6; 12–15^ Extreme cases of cxSVs, such as chromothripsis, have also been identified in both cancer cells and the germline, and involve hundreds of rearrangements often concerning more than one chromosome.^16; 17 11^

Nevertheless, the study of cxSVs has been challenging due to technical limitations. Complex SVs have been reported in projects such as the 1000 Genomes,^1; 18^ but these primarily focused on the canonical types.^19–21^ With the rapid expansion of high-throughput sequencing (HTS) technologies including long-read WGS, genome-wide characterization of SVs with high precision has been achieved,^1^ facilitating the study of more complex forms of SVs. However, systematic identification of Mendelian disease-causing cxSVs in large cohorts is lacking. Here we aim to systematically investigate the role that cxSVs may play in rare Mendelian disorders in our cohort of 1,324 individuals using short-read and long-read WGS.

All participants were consented for the NIHR BioResource research study, which performs WGS of individuals with undiagnosed rare disorders. They were recruited from three different sub-projects: 725 were in the Inherited Retinal Disorders (IRD) project, 472 were in the Neurological and Developmental Disorders (NDD) project, and 127 were in the Next Generation Children (NGC) project, which performs diagnostic trio WGS of individuals from Neonatal and Paediatric Intensive Care Units. All participants provided written informed consent. The study was approved by the East of England Cambridge South national institutional review board (13/EE/0325).

We performed short-read WGS on all the subjects as previously described.^22^ To identify cxSVs we followed a three step framework as previously suggested by Quinlan et al.^4^ First we called canonical structural variants from the short-read WGS data, second we identified putative cxSVs by filtering and clustering adjacent SV calls, and third we reconstructed the likely architecture of the variant by a combination of manual analysis of sequencing reads, Sanger sequence analysis of PCR products across the breakpoint junctions, microarrays, and long-read WGS (Figure S1).

Canonical SVs were called by Canvas,^23^ which identifies copy number gains and losses based on read depth, and Manta,^24^ which calls translocations, deletions, tandem duplications, insertions, and inversions, and is based on both paired read fragment spanning and split read evidence. Filtering and clustering these SV calls then revealed putative cxSVs. Briefly, SVs were initially filtered to keep only those that pass standard Illumina quality filters, do not overlap previously reported CNVs in healthy cohorts,^25^ and are rare (Minor Allele Frequency < 0.01 in the NIHR BioResource (n= 9,453)). To cluster the filtered SVs we identified those where at least one breakpoint was within 1Kb of the breakpoint of another SV in the same individual, and categorized them into different types using a custom script implemented in R. We kept only those clusters that overlapped a coding exon of a disease-associated gene. Disease-associated gene lists were assembled from various sources including OMIM, RetNet and literature searches, then curated to ensure they comply with previously described criteria.^26^ The lists comprise 1,423 genes (NDD) and 248 genes (IRD). For NGC participants, trio analysis focused on *de novo* and rare biallelic variant discovery unrestricted by a gene list. We also filtered out putative cxSVs that only included multiple deletions, as these appear to be caused by insertion of a reverse transcribed mature mRNA molecule elsewhere in the genome.^27^ This resulted in 81 candidate cxSVs.

We performed careful inspection of each candidate cxSV using Integrative Genomics Viewer (IGV).^28^ We excluded a proportion that were actually canonical SVs - for example, dispersed duplications are frequently miscalled as an overlapping deletion and tandem duplication ^29^ - leaving 46 likely real cxSVs. Of these, 42 were deemed unlikely to be pathogenic because the participant’s phenotype was not consistent with the phenotype associated with the gene, or they were heterozygous in a gene known to cause disease with a recessive mode of inheritance, with no second hit. To identify second hits within a gene, we considered SNVs and indels filtered for rare, high-quality, coding variants in a disease-associated gene as previously described.^22^ We also excluded cases where pathogenic SNVs or indels in a disease-associated gene could entirely account for the disease.

The four clinically relevant cxSVs were found in four individuals; P1 and P4 from the NGC project, P2 from NDD and P3 from the IRD project. None of them were observed in our internal cohort of 9,453 genomes, ClinVar or DECIPHER. All predicted novel breakpoint junctions were confirmed in all four participants by Sanger sequencing using standard protocols. To confirm predicted copy number changes and regions of homozygosity we used Illumina SNP genotyping array as previously described,^22^ and/or CytoScan^®^ 750K Cytogenetics Solution microarray (Affymetrix). Here we describe the phenotypes and genetic architecture of the four cxSVs.

Participant 1 (P1) presents a *de novo* duplication-inversion-inversion-deletion encompassing *ARID1B* (MIM: 135900) that causes Coffin-Siris syndrome (CSS [MIM: 135900]). This individual was a four-month old female who was born prematurely and presented with characteristic features of CSS as a neonate. CSS is a multiple malformation syndrome characterized by intellectual disability, severe speech impairment, coarse facial features, microcephaly, developmental delay and hypoplastic nails of the fifth digits.^30^

A large cxSV was identified on chromosome 6, comprising a 3.3Mb duplication, two inversions of 4.9Kb and 3.3Mb, and a 16.3Mb deletion (Figure 1A). A total of 87 proteincoding genes were within the structural variant boundaries (Table S1), of which 21 have been previously described as disease-associated in OMIM. The 16.3Mb deletion contains 72 genes, of which only 6 have been reported as associated with autosomal dominant disease or constrained for loss-of-function (LOF) variation in ExAC^31^ (Table S1). Of these 6, only *ARID1B* has been previously reported as disease-associated with a LOF mechanism. Haploinsufficiency of *ARID1B* causes CSS; consistent with the phenotype of P1. We also looked at the 10 autosomal recessive genes within the deletion and did not find a second likely pathogenic variant in any. No disease-associated gene that was present within the duplicated region had been reported to be triplosensitive. Furthermore, the first inversion and the 3’breakpoint of the second inversion were within *CNKSR3* (MIM: 617476). However, *CNKSR3* has not previously been associated with disease, and is not constrained for LOF variation in ExAC, thus the effect of this inversion on the phenotype remains unknown.

**Figure 1.**
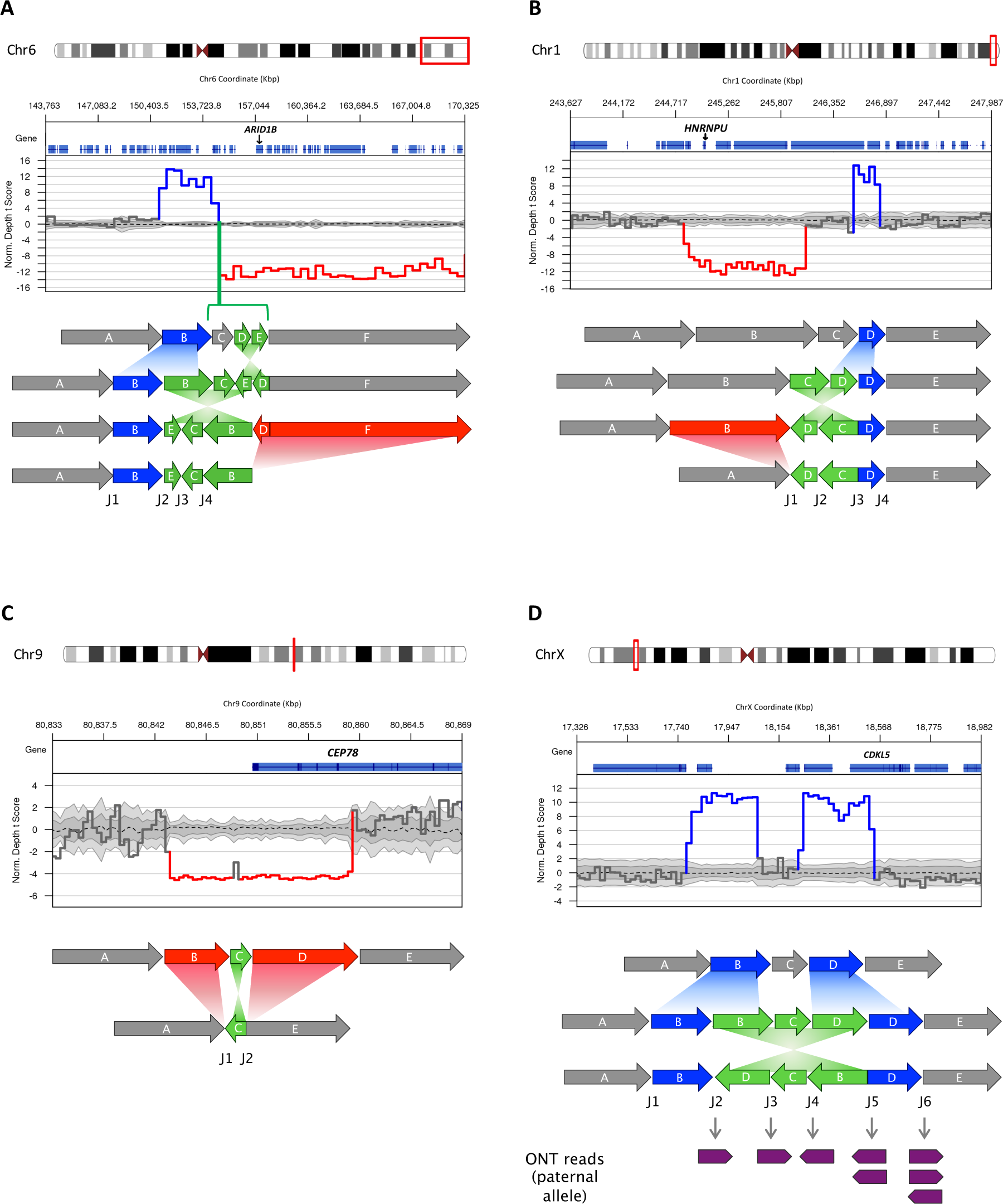
Four Complex Structural Variants Identified by Whole Genome Sequencing. Plots show changes in coverage of short-read WGS (Normalized Depth t Score using CNView, n=250).^51^ Schematic models show the probable sequences of mutational events leading to the formation of the confirmed cxSVs, including putative intermediate derivative chromosomes where relevant. Sizes of fragments are not necessarily to scale. **(A)** A duplication-inversion-inversion-deletion causes Coffin-Siris syndrome in P1. **(B)** A deletion-inversion-duplication causes epilepsy and intellectual disability in P2. **(C)** A deletion-inversion-deletion causes cone-rod dystrophy in P3. **(D)** A duplication-inversion-duplication overlaps with *CDKL5* in P4, who had neonatal hypoxic-ischaemic encephalopathy. Oxford Nanopore Technology (ONT) long-read WGS confirms the presence of a disrupted (J2) and intact (J6) copy of the gene. Only paternally-inherited reads overlapping the junction breakpoints are shown.

For P1, Sanger sequencing of the PCR product generated over the breakpoint junctions confirmed J2 and J3 (Figure 1A). Microarray was performed concurrently with WGS, which confirmed the copy number changes. We observed that the duplication-inversion-inversion-deletion variant occurred on the paternal chromosome, by looking at the parental origin of the hemizygous variants in the deleted region. Therefore, we found a *de novo* cxSV (dupINVinvDEL) in an individual with CSS, classified as pathogenic according to the ACMG guidelines.^32^ Although the LOF of *ARID1B* likely explains the phenotype of this individual, she presented with a very severe case of CSS, so it is possible that other genes affected by the cxSV might contribute to the phenotype.

Participant 2 (P2) has a deletion-inversion-duplication encompassing *HNRNPU* (MIM: 602869). This individual is a 22-year-old male who presented at term with hypotonia. All his early developmental milestones were delayed and he presented with tonic-clonic seizures at 9 months. His seizure disorder has been managed by medication but has continued episodically into adulthood. He presented with significant intellectual disability, autism, and limited speech and language, and MRI showed partial agenesis of the corpus callosum and enlarged ventricles. We identified a cxSV on chromosome 1, formed by a 1.2 Mb deletion and a 246Kb duplication flanking an inversion of 505Kb (Figure 1B). This variant encompassed 8 genes (Table S1), of which two were previously associated with disease: *COX20* (MIM: 614698) and *HNRNPU*, both within the deletion boundaries. Haploinsufficiency of *COX20* was not deemed likely to be pathogenic as variants in this gene have an autosomal recessive mode of inheritance and result in a mitochondrial complex IV deficiency (MIM: 220110), which is not consistent with the individual’s phenotype and no second rare variant was identified. However, *HNRNPU* is a highly constrained gene for LOF variants, in which haploinsufficiency causes early infantile epileptic encephalopathy (EIEE [MIM: 617391]). Microdeletions of *HNRNPU* have been described in individuals with intellectual disability and other clinical features, such as seizures, corpus callosum abnormalities and microcephaly.^33^ The novel genomic junctions generated by the deletion-inversion-duplication (J1 and J3, Figure1B) were confirmed by Sanger sequence analysis, and the deletion and duplication were also confirmed by Illumina SNP genotyping array. Thus, the variant was deemed to be likely pathogenic and is assumed to be *de novo.* Parental segregation is in progress.

Participant 3 (P3), a 66-year-old male, presented with a cone-rod dystrophy and hearing loss due to a homozygous deletion-inversion-deletion overlapping *CEP78* (MIM: 617110). Onset was in his fifth decade with central vision loss, photophobia and nystagmus accompanied by progressive hearing impairment, following a severe influenza-like viral infection. Two homozygous deletions in chromosome 9 of nearly 6 and 10 Kb were found flanking an inversion of 298bp (Figure 1C). The second deletion intersects with the first 5 exons of *CEP78*. Biallelic LOF variants in this gene have been previously shown to cone-rod dystrophy and hearing loss (MIM: 617236).^34; 35^ The new formed junctions J1 and J2 (Figure 1C) were confirmed by Sanger sequencing and both deletions were also confirmed by Illumina SNP genotyping array, although we could not perform segregation analysis due to lack of parental DNA. It was observed to be within a copy number neutral region of homozygosity (ROH), that comprised approximately Chr9:70984372-86933884. This variant was classified as pathogenic.

Participant 4 (P4) presents a duplication-inversion-duplication overlapping *CDKL5* (MIM: 300203) on chromosome X. This individual was a female term (41+1) neonate who presented with foetal bradycardia. She was diagnosed with hypoxic-ischemic encephalopathy (HIE) grade 2, intrauterine hypoxia, and perinatal asphyxia, with poor cord gases. Hypothermia was induced after birth for 72 hours to reduce brain injury. WGS revealed a *de novo* duplication-inversion-duplication, with the respective sizes of 280Kb, 458Kb, and 283Kb (Figure 1D). The inversion 3’ breakpoint is in intron 4 of 21 of *CDKL5* (NM_001037343) (Figure 1D). Heterozygous rare variants in X-linked *CDKL5* in females cause Early Infantile Epileptic Encephalopathy (EIEE), severe intellectual disability and Rett-like features (MIM: 300672). There are three other genes within the boundaries of this cxSV, none of them disease-associated in OMIM.

The Manta and Canvas calls from short-read WGS suggested two possible molecular models. Model one: formed by two closely located duplications followed by an inversion affecting only one copy of each duplicated segment (Figure S2A).^12^ And model two: formed by two closely located duplications followed by an inversion affecting both copies of the 3’ duplicated segment (Figure S2B). In both models the non-duplicated nested fragment (named C) is inverted. Importantly, model one suggests that there is an intact copy of *CDKL5* on the non-reference allele, in addition to the disrupted copy; model two does not. Therefore, resolving the model was essential for the interpretation of the pathogenicity of this variant. We attempted PCR amplification over the predicted new formed junctions, and only the one supporting the disrupted *CDKL5*, common to both, was amplified and confirmed by Sanger sequencing (Figure2S). Both duplications were confirmed by microarray.

In order to resolve the model, we performed long-read WGS of DNA from this individual using Oxford Nanopore Technologies (ONT). The sample was prepared using the 1D ligation library prep kit (SQK-LSK108), and genomic libraries were sequenced on R9 flowcell. Read sequences were extracted from base-called FAST5 files by albacore (version 2.0.2) to generate FASTQ files, and then aligned against the GRCh37 human reference genome using NGMLR (version 0.2.6)^36^ and LAST (version 912).^37^ Analysis was performed using default parameters, and for LAST we used first last-train function to optimise alignment scoring.

We obtained a median read length of 8,136bp (Figure S3A), 56% of the genome was covered with a minimum coverage of 3x (Figure S3B), and around 97% of the reads mapped to the human genome (GRCh37). All the breakpoints of the cxSV were covered by at least four reads.

Coverage was insufficient to resolve the cxSV using long-read SV calling algorithms such as Sniffles^36^ or NanoSV,^38^ (for which a minimum coverage of 10x is recommended). In lieu of this, we manually reviewed the split long reads across the cxSV junction breakpoints. Eight of the reads that covered the cxSV breakpoints were identified as inherited from the paternal chromosome, either by SNP phasing (Figure 1D, J2, J3, J4 and J6) or by indirect phasing based on the assumption that breakpoint junctions occur on the same allele (Figure 1D, J5). Therefore, ONT sequencing allowed us to identify two reads supporting the junction that was initially not possible to confirm by Sanger sequencing (J5) due to sequence homology. By phasing analysis, we were also able to identify three reads supporting an intact copy of *CDKL5* in the allele inherited from the father, confirming that model one (Figure 1D) is likely to be correct. Although we were able to confirm that the cxSV harbours an intact copy of *CDKL5*, we cannot conclude that the *de novo* cxSV identified is benign and unrelated to the neonatal phenotype since we cannot rule out the possibility that regulatory elements are affected. Thus, this variant was classified as a variant of unknown significance (VUS). The child is currently one year old and developmentally normal with no seizures, but remains under ongoing follow up.

It has previously been reported that DNA replication-based mechanisms such as microhomology-mediated break-induced replication (MMBIR) or fork stalling and template switching (FoSTeS) are likely to be the primary mechanism responsible for the formation of cxSVs.^4; 5; 39–41^ Our data overall support this as there is microhomology of at least 3bp in 8 of the 9 novel breakpoint junctions in the four individuals (Table S2). We also observe in P4 the insertion of a 100bp *Alu* sequence in J2 junction. It has been previously suggested that *Alu* elements could facilitate template switching and annealing via microhomology between replication forks.^41^ However, additional evaluation of the breakpoint sequences with RepeatMasker also identified longer repeats and low complexity sequences in three of the individuals. In P1 we found that sequence flanking two of the breakpoints had high homology with SINE sequences (ERVL-MaLRs), one with LINE sequences (L2) and one with DNA/hAT-Charlie (MER3) sequences (Table 1). In P2 we noted that sequence flanking three of the breakpoints had homology with SINE sequences (*Alu* and MIR), and in P3, sequences surrounding all the breakpoints presented high homology with LINEs. This raises the possibility of involvement of homologous recombination mechanisms.^5^ Interestingly, we also note that the inversions in P1, P2 and P4 involved junctions that were newly formed by the duplications in our predicted models.

**Table 1.**
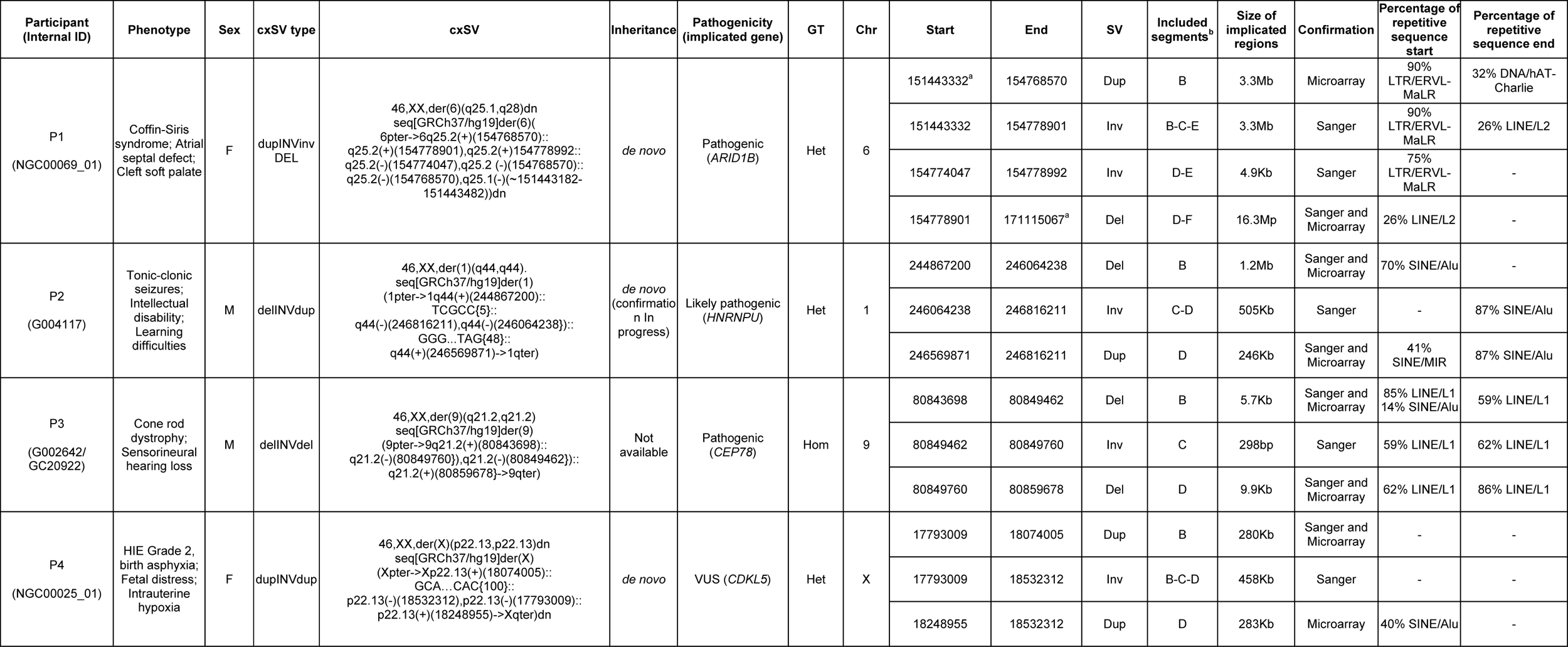
Complex Structural Variant Characteristics. cxSVs are described using Next-Gen Cytogenetic Nomenclature.^52^ For details of all affected genes see Table S1. ^a^Not confirmed by Sanger sequencing; coordinate obtained from direct observation of WGS data in IGV. ^b^Refers to genomic segments as shown in Figure 1. Percentage of repetitive sequence calculated using RepeatMasker for a sequence of 300bp across the breakpoint. cxSV=complex structural variant, GT=genotype, Chr=chromosome, SV=structural variant.

| Participant<br>(Internal ID) | Phenotype | Sex | cxSV type | cxSV | Inheritance | Pathogenicity<br>(implicated gene) | GT | Chr | Start | End | SV | Included<br>segments<sup>b</sup> | Size of<br>implicated<br>regions | Confirmation | Percentage of<br>repetitive<br>sequence<br>start | Percentage of<br>repetitive<br>sequence<br>end |
| --- | --- | --- | --- | --- | --- | --- | --- | --- | --- | --- | --- | --- | --- | --- | --- | --- |
| P1<br>(NGC00069_01) | Coffin-Siris<br>syndrome; Atrial<br>septal defect;<br>Cleft soft palate | F | dupINVinv<br>DEL | 46,XX,der(6)(q25.1.q28)dn<br>seq[GRCh37/hg19]der(6)(<br>6pter->6q25.2(+)(154768570)::<br>q25.2(+)(154778901),q25.2(+)(154778992::<br>q25.2(-)(154774047),q25.2(-)(154768570)::<br>q25.2(-)(154768570),q25.1(-)(~151443182-151443482))dn | *de novo* | Pathogenic<br>(ARID1B) | Het | 6 | 151443332<sup>a</sup> | 154768570 | Dup | B | 3.3Mb | Microarray | 90%<br>LTR/ERVL-MaLR | 32% DNA/hAT-Charlie |
|  |  |  |  |  |  |  |  |  | 151443332 | 154778901 | Inv | B-C-E | 3.3Mb | Sanger | 90%<br>LTR/ERVL-MaLR | 26% LINE/L2 |
|  |  |  |  |  |  |  |  |  | 154774047 | 154778992 | Inv | D-E | 4.9Kb | Sanger | 75%<br>LTR/ERVL-MaLR | - |
|  |  |  |  |  |  |  |  |  | 154778901 | 171115067<sup>a</sup> | Del | D-F | 16.3Mp | Sanger and<br>Microarray | 26% LINE/L2 | - |
| P2<br>(G004117) | Tonic-clonic<br>seizures;<br>Intellectual<br>disability;<br>Learning<br>difficulties | M | delINVdup | 46,XX,der(1)(q44.q44).<br>seq[GRCh37/hg19]der(1)<br>(1pter->1q44(+)(244867200)::<br>TCGCC[5];<br>q44(-)(246816211),q44(-)(246064238)::<br>GGG...TAG[48];<br>q44(+)(246569871)->1qter) | *de novo*<br>(confirmatio<br>n In<br>progress) | Likely pathogenic<br>(HNRNPU) | Het | 1 | 244867200 | 246064238 | Del | B | 1.2Mb | Sanger and<br>Microarray | 70% SINE/Alu | - |
|  |  |  |  |  |  |  |  |  | 246064238 | 246816211 | Inv | C-D | 505Kb | Sanger | - | 87% SINE/Alu |
|  |  |  |  |  |  |  |  |  | 246569871 | 246816211 | Dup | D | 246Kb | Sanger and<br>Microarray | 41%<br>SINE/MIR | 87% SINE/Alu |
| P3<br>(G002642/<br>GC20922) | Cone rod<br>dystrophy;<br>Sensorineural<br>hearing loss | M | delINVdel | 46,XX,der(9)(q21.2.q21.2)<br>seq[GRCh37/hg19]der(9)<br>(9pter->9q21.2(+)(80843698)::<br>q21.2(-)(80849760)),q21.2(-)(80849462)::<br>q21.2(+)(80859678)->9qter) | Not<br>available | Pathogenic<br>(CEP78) | Hom | 9 | 80843698 | 80849462 | Del | B | 5.7Kb | Sanger and<br>Microarray | 85% LINE/L1<br>14% SINE/Alu | 59% LINE/L1 |
|  |  |  |  |  |  |  |  |  | 80849462 | 80849760 | Inv | C | 298bp | Sanger | 59% LINE/L1 | 62% LINE/L1 |
|  |  |  |  |  |  |  |  |  | 80849760 | 80859678 | Del | D | 9.9Kb | Sanger and<br>Microarray | 62% LINE/L1 | 86% LINE/L1 |
| P4<br>(NGC00025_01) | HIE Grade 2,<br>birth asphyxia;<br>Fetal distress;<br>Intrauterine<br>hypoxia | F | dupINVdup | 46,XX,der(X)(p22.13,p22.13)dn<br>seq[GRCh37/hg19]der(X)<br>(Xpter->Xp22.13(+)(18074005)::<br>GCA...CAC[100];<br>p22.13(-)(18532312),p22.13(-)(17793009)::<br>p22.13(+)(18248955)->Xqter)dn | *de novo* | VUS (CDKL5) | Het | X | 17793009 | 18074005 | Dup | B | 280Kb | Sanger and<br>Microarray | - | - |
|  |  |  |  |  |  |  |  |  | 17793009 | 18532312 | Inv | B-C-D | 458Kb | Sanger | - | - |
|  |  |  |  |  |  |  |  |  | 18248955 | 18532312 | Dup | D | 283Kb | Microarray | 40% SINE/Alu | - |

In this study, we evaluate the role of cxSVs in Mendelian disorders using short-read and long-read WGS. By analysing cxSVs from WGS data in 1,324 individuals with NDD or IRD, we found that 46 (3.5%) have a rare cxSVs overlapping disease-associated genes, and at least 3 (0.3%) are disease-causing. This compares to approximately 5-20% of individuals with Mendelian disorders who have a clinically relevant canonical SV.^22; 42; 43^ However, 0.3% is likely to be an underestimate due to three reasons.

First, a substantial proportion of individuals in NDD and IRD cohorts are pre-screened by microarray prior to enrolment in the project. It has been reported that 7.6% of all rare duplications detected by microarray are part of a complex rearrangement.^12^ This misclassification is in part due to the impossibility for detecting inversions by microarray, since inversions are involved in 84.8% of cxSVs.^11^ Thus, it is likely that many ‘canonical CNVs’ detected by microarray are actually misclassified cxSVs.

Second, short-read WGS is limited in identifying and resolving these variants, since they are prone to occur at repetitive regions where mapping and variant calling algorithms have lower sensitivity. Certain categories of cxSVs such as chromothripsis and those formed by nested rather than chained breakpoints could also be excluded by our filtering and clustering method. These limitations may now be addressed by long-read sequencing technologies such as Nanopore, which have the advantage of reads of 10-100Kb allowing for more accurate mapping particularly over repetitive regions, and facilitating phasing.^38^ Although coverage is lower and error rate is higher than short-read WGS, various studies have demonstrated the power of long-read WGS to detect SVs and cxSVs.^36; 38; 44–46^

Third, estimating the frequency of cxSVs requires defining them, which is not always straightforward because the distinction between canonical SVs, cxSVs, and chromoanagenesis can be unclear.^39; 47^ It is therefore perhaps appropriate to consider types of human genomic variation as a continuum rather than discrete classes, progressing from SNVs (that cause the least disruption to the genome), through indels, canonical SVs and cxSVs to the highly disruptive chromoanagenesis and aneuploidies.

The low frequency of pathogenic cxSVs compared to other classes of pathogenic variants is likely to be due to differential rates of the causative mutational processes and higher purifying selection pressure on cxSVs.^2; 11^ Consequently, it is unsurprising that three of the cxSVs reported here (in P1, P2 (confirmation in progress), and P4) are *de novo*. Two were also confirmed to have occurred on the paternally inherited allele, consistent with previously reported observations that ∼80% of *de novo* mutations are of paternal origin.^48; 49^

Complex SVs are not typically considered in research and clinical diagnostic pipelines, in part due to the technical and analytical challenges around identification and interpretation. Therefore, when analysis is restricted to canonical SVs alone, the full scope of genome-wide structural variation is overlooked. This study demonstrates the potential for detecting and characterising cxSVs from short-read WGS to identify possible clinically relevant cxSVs, then performing long-read sequencing where necessary to resolve breakpoints. An alternative possible method is long insert WGS (liWGS), which has been successfully employed to detect cxSVs in other studies, and has the advantage of improved mapping particularly over repetitive regions due to the large fragments, but has a lower resolution of ∼5Kb. ^11–13^

Our experience with P4, who has a VUS involving *CDKL5*, demonstrates that understanding the genetic architecture of a cxSV is essential for interpreting the pathogenicity of the variant, especially if the gene of interest is disrupted by a duplication or inversion rather than a deletion. The impact of a deletion on the function of affected genes is generally assumed to be LOF. However, the consequence of a duplication can be uncertain and depends on precisely how the variant rearranges the gene, as well as gene-specific factors such as dosage sensitivity. Furthermore, duplications intersecting regulatory regions can result in a different phenotype from variants within the gene itself.^50^

Investigating cxSVs in our cohort identified previously reported subclasses (delINVdup, delINVdel and dupINVdup in P2, P3 and P4 respectively), as well as a dupINVinvDEL in P1.^11^ As has been previously anticipated, with the study of larger population-scale cohorts and the use of higher resolution technologies such as long-read WGS, many additional subclasses will continue to be discovered.^11^ However, inferring the likely molecular mechanism of cxSV formation is challenging due to the complexity of the events. The high frequency of microhomology observed at the breakpoint junctions of the cxSVs in our participants and the presence of an inserted sequence in three of them is generally consistent with the hypothesis that replication-based mechanisms such as FoSTeS/MMBIR are the primary mechanism responsible for the formation of cxSVs. However, we also find evidence for variable number tandem repeat (VNTR) sequences in all the participants. Taken together, these results suggest a hypothesis whereby multiple mutational mechanisms could mediate the formation of cxSVs, as novel sequence junctions in intermediate derivative chromosomes formed by canonical SV could facilitate additional mutational events.

Our work demonstrates that cxSVs contribute to rare Mendelian disorders and provides insight into identifying and resolving cxSVs by using both short and long-read WGS. We suggest that cxSVs should be included into research and clinical diagnosis and considered when screening SVs in the human genome. Detailed characterization of cxSVs and large-scale WGS studies are essential for unveiling the complex architecture and determining accurate population frequencies. Furthermore, we demonstrate a combinatorial approach of short-read WGS and confirmation by microarray, Sanger sequencing, and long-read WGS as an approach for resolving and characterizing cxSVs.

## Supplemental data

Supplemental data includes three figures and two tables.

## Consortia

### NIHR BioResource Principal Investigators

Tim Aitman, David Bennett, Mark Caulfield, Patrick F Chinnery, Peter Dixon, Daniel Gale, Ania Koziell, Taco Kuijpers, Michael Laffan, Eamonn Maher, Hugh S Markus, Nicholas Morrell, Willem Ouwehand, F Lucy Raymond, Irene Roberts, Kenneth G C Smith, James Thaventhiran, Adrian Thrasher, Hugh Watkins, Catherine Williamson, Geoff Woods

### Ethics, Governance, Recruitment coordination and Clinical Bioinformatics

Sofie Ashford, Matthew Brown, Naomi Clements Brod, Eleanor Dewhurst, Marie Erwood, Amy Frary, Csaba Halmagyi, Roger James, Rachel Linger, Jennifer Martin, Sofia Papadia, Crina Samarghitean, Emily Staples, Hannah Stark, Catherine Titterton, Julie von Ziegenweidt, Katherine Yates, Ping Yu

### Sample and Data processing: EMBL-European Bioinformatics Institute

Jeff Almeida-King; **GENALICE** Jack Findhammer, Johannes Karten, Tim Karten, Bas Tolhuis, Maarten Vandekuilen; **High Performance Computing Facility** Paul Calleja, Robert Klima, Stuart Rankin, Wojciech Turek; **Illumina** David Bentley, Christian Bourne, Camilla Colombo, Claire Geoghegan, Terry Gerighty, Russell Grocock, Joseph Hughes, Sean Humphray, Sarah Hunter, John Peden, Christine Rees; **University of Cambridge** Anthony Attwood, Christine Bryson, Abigail Crisp-Hihn, Sri V V Deevi, Karen Edwards, James Fox, Fengyuan Hu, Jennifer Jolley, Rutendo Mapeta, Stuart Meacham, Christopher Penkett, Paula Rayner-Matthews, Olga Sharmardina, Ilenia Simeoni, Simon Staines, Jonathan Stephens, Kathleen Stirrups, Salih Tuna, Christopher Watt, Deborah Whitehorn, Yvette Wood

### Method Development: High Performance Computing Facility

Stuart Rankin; **MRC Biostatistics Unit** Henry Farmery, Sylvia Richardson; **University of Cambridge** Keren Carss, Stefan Gräf, Daniel Greene, Matthias Haimel, Ernest Turro; **Wellcome Trust Sanger Institute** Patrick Short

### Data Analysis

Micheala Aldred, William Astle, Chris Babbs, Agnieszka Bierzynska, Marta Bleda, Oliver Burren, Keren Carss, Courtney French, Kathleen Freson, Pavandeep Ghataorhe, Stefan Gräf, Daniel Greene, Charaka Hadinnapola, Matthias Haimel, Joshua Hodgson, Ania Koziell, Hana Lango Allen, Adam Levine, Wei Li, Bin Liu, Eleni Louka, Adam Mead, Karyn Megy, Monika Mozere, Jennifer O’Sullivan, Steven Okoli, David Parry, Inga Prokopenko, Beth Psaila, Anupama Rao, Augusto Rendon, Christopher J Rhodes, Irene Roberts, Noemi Roy, Omid Sadeghi-Alavijeh, Alba Sanchis-Juan, Katherine Smith, Mark Southwood, Daniel Stein, Emilia Swietlik, Rhea Tan, James Thaventhiran, Ernest Turro, Paul Upton, Natalie Van Zuydam, Wei Wei, James Whitworth

### Clinical Interpretation and Multi-Disciplinary Teams

Stephen Abbs, Tim Aitman, Philip Ancliff, Gavin Arno, Chiara Bacchelli, David Bennett, Agnieszka Bierzynska, Siobhan Burns, Keren Carss, Louise Daugherty, Peter Dixon, Kate Downes, Anna Drazyk, Courtney French, Kathleen Freson, Daniel Gale, Kimberley Gilmour, Keith Gomez, Stefan Gräf, Detelina Grozeva, Charaka Hadinnapola, Simon Holden, Ania Koziell, Taco Kuijpers, Michael Laffan, Hana Lango Allen, Mark Layton, Adam Levine, Eleni Louka, Eamonn Maher, Jesmeen Maimaris, Rutendo Mapeta, Hugh S Markus, Jennifer Martin, Karyn Megy, Sarju Mehta, Nicholas Morrell, Andrew Mumford, Willem Ouwehand, David Parry, F Lucy Raymond, Irene Roberts, Noemi Roy, Moin Saleem, Alba Sanchis-Juan, Sinisa Savic, Ilenia Simeoni, Emilia Swietlik, Rhea Tan, James Thaventhiran, Andreas Themistocleous, David Thomas, Marc Tischkowitz, Matthew Traylor, Ernest Turro, Natalie Van Zuydam, Anthony Vandersteen, Andrew Webster, James Whitworth, Catherine Williamson, Geoff Woods

### Steering groups

**NIHR BioResource Sequencing and Informatics Committee (SIC)** Gerome Breen, John Chambers, Matthew Hurles, Nathalie Kingston, Mark McCarthy, Willem Ouwehand, F Lucy Raymond, Nilesh Samani, Michael Simpson, Nicholas Wood; **NIHR BioResource – Rare Diseases Senior Management Team (SMT)** Sofie Ashford, John Bradley, Debra Fletcher, Roger James, Mary Kasanicki, Nathalie Kingston, Willem Ouwehand, Christopher Penkett, F Lucy Raymond, Hannah Stark, Kathleen Stirrups, Tim Young

### Rare Disease and Control Collections

**Bleeding, thrombotic and Platelet Disorders (BPD)** David Allsup, Tamam Bakchoul, Tadbir Bariana, Tina Biss, Sara Boyce, Janine Collins, Peter Collins, Nicola Curry, Kate Downes, Tina Dutt, Wendy Erber, Gillian Evans, Tamara Everington, Remi Favier, Kathleen Freson, Keith Gomez, Daniel Greene, Andreas Greinacher, Paolo Gresele, Daniel Hart, Rashid Kazmi, Michael Laffan, Michele Lambert, Claire Lentaigne, Bella Madan, Sarah Mangles, Mary Mathias, Carolyn Millar, Andrew Mumford, Samya Obaji, Willem Ouwehand, Sofia Papadia, Kathelijne Peerlinck, Catherine Roughley, Sol Schulman, Marie Scully, Susie Shapiro, Keith Sibson, Suthesh Sivapalaratnam, R Campbell Tait, Kate Talks, Chantal Thys, Cheng-Hock Toh, Ernest Turro, Chris Van Geet, Sarah Westbury, John-Paul Westwood; **Cerebral Small Vessel Disease (CSVD)** Rhea Tan, Julian Barwell, Kate Downes, Stefan Gräf, Kirsty Harkness, Sarju Mehta, Keith Muir, Ahamad Hassan, Matthew Traylor, Anna Drazyk, Hugh S Markus; **Ehlers Danlos Syndrome (EDS)** David Parry, Munaza Ahmed, Alex Henderson, Hanadi Kazkaz, Anthony Vandersteen, Tim Aitman; **Hypertrophic Cardiomyopathy (HCM)** Elizabeth Ormondroyd, Kate Thomson, Timothy Dent, Paul Brennan, Rachel Buchan, Teofila Bueser, Gerald Carr-White, Stuart Cook, Matthew Daniels, Andrew Harper, Alex Henderson, James Ware, Hugh Watkins; **Intrahepatic Cholestasis of Pregnancy (ICP)** Peter Dixon, Jennifer Chambers, Floria Cheng, Maria C Estiu, William Hague, Hanns-Ulrich Marschall, Marta Vazquez-Lopez, Catherine Williamson; **Inherited Retinal Disorders (IRD)** Gavin Arno, Keren Carss, Eleanor Dewhurst, Marie Erwood, Courtney French, Michel Michaelides, Tony Moore, F Lucy Raymond, Alba Sanchis-Juan, Andrew Webster; **Leber Hereditary Optic Neuropathy (LHON)** Patrick F Chinnery, Philip Griffiths, Rita Horvath, Gavin Hudson, Neringa Jurkute, Angela Pyle, Wei Wei, Patrick Yu-Wai-Man; **Multiple Primary Malignant Tumours (MPMT)** James Whitworth, Julian Adlard, Munaza Ahmed, Ruth Armstrong, Julian Barwell, Carole Brewer, Ruth Casey, Trevor Cole, Dafydd Gareth Evans, Lynn Greenhalgh, Helen Hanson, Alex Henderson, Jonathan Hoffmann, Louise Izatt, Ajith Kumar, Fiona Lalloo, Kai Ren Ong, Soo-Mi Park, Joan Paterson, Claire Searle, Lucy Side, Katie Snape, Emma Woodward, Marc Tischkowitz, Eamonn Maher; **Neurological and Developmental Disorders (NDD)** Keren Carss, Eleanor Dewhurst, Marie Erwood, Courtney French, Detelina Grozeva, Manju Kurian, F Lucy Raymond, Alba Sanchis-Juan; **Neuropathic Pain Disorders (NPD)** Andreas Themistocleous, David Gosal, Rita Horvath, Andrew Marshall, Emma Matthews, Mark McCarthy, Tara Renton, Andrew Rice, Tom Vale, Natalie Van Zuydam, Suellen Walker, Geoff Woods, David Bennett; **Primary Immune Disorders (PID)** Hana Alachkar, Richard Antrobus, Michael Browning, Matthew Buckland, Siobhan Burns, Oliver Burren, Nichola Cooper, Elizabeth Drewe, David Edgar, William Egner, Kimberley Gilmour, Sarah Goddard, Pavels Gordins, Sofia Grigoriadou, Scott Hackett, Grant Hayman, Aarnoud Huissoon, Stephen Jolles, Peter Kelleher, Taco Kuijpers, Dinakantha Kumararatne, Hana Lango Allen, Hilary Longhurst, Jesmeen Maimaris, Alex Richter, Ravishankar Sargur, Sinisa Savic, Carrock Sewell, Ilenia Simeoni, Kenneth G C Smith, Emily Staples, James Thaventhiran, Moira Thomas, David Thomas, Adrian Thrasher, Steven Welch, Austen Worth, Patrick Yong; **Primary Membranoproliferative Glomerulonephritis (PMG)** Adam Levine, Omid Sadeghi-Alavijeh, Edwin Wong, Terry Cook, Martin Christian, Matthew Hall, Claire Harris, Paul McAlinden, Kevin Marchbank, Stephen Marks, Heather Maxwell, Monika Mozere, Julie Wessels, MPGN/C3 Glomerulopathy Rare Renal Disease group, Sally Johnson, Daniel Gale; **Pulmonary Arterial Hypertension (PAH)** Marta Bleda, Harm Bogaard, Colin Church, Gerry Coghlan, Robin Condliffe, Paul Corris, Cesare Danesino, Mélanie Eyries, Henning Gall, Stefano Ghio, Hossein-Ardeschir Ghofrani, Simon Gibbs, Barbara Girerd, Stefan Gräf, Charaka Hadinnapola, Matthias Haimel, Simon Holden, Arjan Houweling, Luke Howard, Marc Humbert, David Kiely, Gabor Kovacs, Allan Lawrie, Robert MacKenzie Ross, Jennifer Martin, Shahin Moledina, David Montani, Nicholas Morrell, Michael Newnham, Andrea Olschewski, Horst Olschewski, Andrew Peacock, Joanna Pepke-Zaba, Laura Scelsi, Werner Seeger, Florent Soubrier, Jay Suntharalingam, Emilia Swietlik, Mark Toshner, Carmen Treacy, Richard Trembath, Anton Vonk Noordegraaf, Quintin Waisfisz, John Wharton, Martin R Wilkins, Stephen Wort, Katherine Yates; **Stem cell and Myeloid Disorders (SMD)** Eleni Louka, Noemi Roy, Anupama Rao, Philip Ancliff, Chris Babbs, Mark Layton, Adam Mead, Jennifer O’Sullivan, Steven Okoli, Irene Roberts; **Steroid Resistant Nephrotic Syndrome (SRNS)** Agnieszka Bierzynska, Elizabeth Colby, Simon Satchell, Moin Saleem, Ania Koziell.

### Next Generation Children (NGC) project

Steve Abbs, Shruti Agrawal, Gautam Ambegaonkar, June Anne Gold, Ruth Armstrong, Topun Austin, Kathryn Beardsall, Gusztav Belteki, Marion Bohatschek, Sarah Bowdin, Susan Broster, Rosalie Campbell, Keren Carss, Rajiv Chaudhary, Manali Chitre, Cristine Costa, Angela D’Amore, Isabelle Delon, Eleanor Dewhurst, Helen Dolling, Carolyn Dunn, Helen Firth, Annie Fitzsimmons, Courtney French, Ricardo Garcia Branco, Jennifer Hague, Joanne Harley, Natalie Holder, Shazia Hoodbhoy, Ian Johnson, Riaz Kayani, Wilf Kelsall, Deepa Krishnakumar, Anna Maw, Karyn Mégy, Sarju Mehta, Roddy O’Donnell, Samantha O’Hare, Amanda Ogilvy-Stuart, Stergios Papakostas, Soo-Mi Park, Alasdair Parker, Nazima Pathan, Chris Penkett, Matina Prapa, Dilna Pushpajan, F Lucy Raymond, David Rowitch, Audrienne Sammut, Alba Sanchis-Juan, Katherine Schon, Olga Shamardina, Yogen Singh, Kelly Spike, Jonathan Stephens, Kathy Stirrups, Analisa Taylor Tavares, Salih Tuna, Doris Wari-Pepple, Geoff Woods

## Acknowledgments

We thank the participants involved in this study and their families. This work was supported by The National Institute for Health Research England (NIHR) for the NIHR BioResource project (grant number RG65966). The NIHR BioResource projects were approved by Research Ethics Committees in the UK and appropriate national ethics authorities in non-UK enrolment centres. We thank Dr Ernest Turro for his part in initiating our collaboration with Oxford Nanopore Technologies, and for helpful comments on this manuscript. GA and ARW are supported by the NIHR Biomedical Research Centre at Moorfields Eye Hospital NHS Trust and UCL Institute of Ophthalmology, Moorfields Eye Charity, Fight for Sight (UK), Foundation Fighting Blindness and Retinitis Pigmentosa Fighting Blindness. GA is a recipient of a Fight for Sight (UK) Early Career Investigator Award.

## Web resources

1. OMIM, http://www.omim.org
2. ExAC, http://exac.broadinstitute.org
3. gnomAD, http://gnomad.broadinstitute.org
4. Ensembl Genome Browser, http://www.ensembl.org/index.html
5. UCSC genome browser, http://genome.ucsc.edu
6. RetNet, http://www.sph.uth.tmc.edu/RetNet
7. ClinVar, http://www.ncbi.nlm.nih.gov/clinvar
8. Decipher, http://decipher.sanger.ac.uk/
9. Repeatmasker, http://www.repeatmasker.org

## References

1. Sudmant, P.H., Rausch, T., Gardner, E.J., Handsaker, R.E., Abyzov, A., Huddleston, J., Zhang, Y., Ye, K., Jun, G., Fritz, M.H., et al. (2015). An integrated map of structural variation in 2,504 human genomes. Nature 526, 75–81.

2. Zhang, F., Gu, W., Hurles, M.E., and Lupski, J.R. (2009). Copy number variation in human health, disease, and evolution. Annu Rev Genomics Hum Genet 10, 451–481.

3. Conrad, D.F., Bird, C., Blackburne, B., Lindsay, S., Mamanova, L., Lee, C., Turner, D.J., and Hurles, M.E. (2010). Mutation spectrum revealed by breakpoint sequencing of human germline CNVs. Nat Genet 42, 385–391.

4. Quinlan, A.R., and Hall, I.M. (2012). Characterizing complex structural variation in germline and somatic genomes. Trends Genet 28, 43–53.

5. Weckselblatt, B., and Rudd, M.K. (2015). Human Structural Variation: Mechanisms of Chromosome Rearrangements. Trends Genet 31, 587–599.

6. Carvalho, C.M., Ramocki, M.B., Pehlivan, D., Franco, L.M., Gonzaga-Jauregui, C., Fang, P., McCall, A., Pivnick, E.K., Hines-Dowell, S., Seaver, L.H., et al. (2011). Inverted genomic segments and complex triplication rearrangements are mediated by inverted repeats in the human genome. Nat Genet 43, 1074–1081.

7. Quelin, C., Spaggiari, E., Khung-Savatovsky, S., Dupont, C., Pasquier, L., Loeuillet, L., Jaillard, S., Lucas, J., Marcorelles, P., Journel, H., et al. (2014). Inversion duplication deletions involving the long arm of chromosome 13: phenotypic description of additional three fetuses and genotype-phenotype correlation. Am J Med Genet A 164A, 2504–2509.

8. Arno, G., Agrawal, S.A., Eblimit, A., Bellingham, J., Xu, M., Wang, F., Chakarova, C., Parfitt, D.A., Lane, A., Burgoyne, T., et al. (2016). Mutations in REEP6 Cause Autosomal-Recessive Retinitis Pigmentosa. Am J Hum Genet 99, 1305–1315.

9. Kuroda, Y., Ohashi, I., Saito, T., Nagai, J., Ida, K., Naruto, T., Wada, T., and Kurosawa, K. (2014). Deletion of UBE3A in brothers with Angelman syndrome at the breakpoint with an inversion at 15q11.2. Am J Med Genet A 164A, 2873–2878.

10. Starr, L.J., Truemper, E.J., Pickering, D.L., Sanger, W.G., and Olney, A.H. (2014). Duplication of 20qter and deletion of 20pter due to paternal pericentric inversion: patient report and review of 20qter duplications. Am J Med Genet A 164A, 2020–2024.

11. Collins, R.L., Brand, H., Redin, C.E., Hanscom, C., Antolik, C., Stone, M.R., Glessner, J.T., Mason, T., Pregno, G., Dorrani, N., et al. (2017). Defining the diverse spectrum of inversions, complex structural variation, and chromothripsis in the morbid human genome. Genome Biol 18, 36.

12. Brand, H., Collins, R.L., Hanscom, C., Rosenfeld, J.A., Pillalamarri, V., Stone, M.R., Kelley, F., Mason, T., Margolin, L., Eggert, S., et al. (2015). Paired-Duplication Signatures Mark Cryptic Inversions and Other Complex Structural Variation. Am J Hum Genet 97, 170–176.

13. Brand, H., Pillalamarri, V., Collins, R.L., Eggert, S., O’Dushlaine, C., Braaten, E.B., Stone, M.R., Chambert, K., Doty, N.D., Hanscom, C., et al. (2014). Cryptic and complex chromosomal aberrations in early-onset neuropsychiatric disorders. Am J Hum Genet 95, 454–461.

14. Tabet, A.C., Verloes, A., Pilorge, M., Delaby, E., Delorme, R., Nygren, G., Devillard, F., Gerard, M., Passemard, S., Heron, D., et al. (2015). Complex nature of apparently balanced chromosomal rearrangements in patients with autism spectrum disorder. Mol Autism 6, 19.

15. Lohmann, K., Redin, C., Tonnies, H., Bressman, S.B., Subero, J.I.M., Wiegers, K., Hinrichs, F., Hellenbroich, Y., Rakovic, A., Raymond, D., et al. (2017). Complex and Dynamic Chromosomal Rearrangements in a Family With Seemingly Non-Mendelian Inheritance of Dopa-Responsive Dystonia. JAMA Neurol 74, 806–812.

16. Stephens, P.J., Greenman, C.D., Fu, B., Yang, F., Bignell, G.R., Mudie, L.J., Pleasance, E.D., Lau, K.W., Beare, D., Stebbings, L.A., et al. (2011). Massive genomic rearrangement acquired in a single catastrophic event during cancer development. Cell 144, 27–40.

17. Forment, J.V., Kaidi, A., and Jackson, S.P. (2012). Chromothripsis and cancer: causes and consequences of chromosome shattering. Nat Rev Cancer 12, 663–670.

18. Hehir-Kwa, J.Y., Marschall, T., Kloosterman, W.P., Francioli, L.C., Baaijens, J.A., Dijkstra, L.J., Abdellaoui, A., Koval, V., Thung, D.T., Wardenaar, R., et al. (2016). A high-quality human reference panel reveals the complexity and distribution of genomic structural variants. Nat Commun 7, 12989.

19. Lander, E.S., Linton, L.M., Birren, B., Nusbaum, C., Zody, M.C., Baldwin, J., Devon, K., Dewar, K., Doyle, M., FitzHugh, W., et al. (2001). Initial sequencing and analysis of the human genome. Nature 409, 860–921.

20. Venter, J.C., Adams, M.D., Myers, E.W., Li, P.W., Mural, R.J., Sutton, G.G., Smith, H.O., Yandell, M., Evans, C.A., Holt, R.A., et al. (2001). The sequence of the human genome. Science 291, 1304–1351.

21. Genomes Project, C., Abecasis, G.R., Auton, A., Brooks, L.D., DePristo, M.A., Durbin, R.M., Handsaker, R.E., Kang, H.M., Marth, G.T., and McVean, G.A. (2012). An integrated map of genetic variation from 1,092 human genomes. Nature 491, 56–65.

22. Carss, K.J., Arno, G., Erwood, M., Stephens, J., Sanchis-Juan, A., Hull, S., Megy, K., Grozeva, D., Dewhurst, E., Malka, S., et al. (2017). Comprehensive Rare Variant Analysis via Whole-Genome Sequencing to Determine the Molecular Pathology of Inherited Retinal Disease. Am J Hum Genet 100, 75–90.

23. Roller, E., Ivakhno, S., Lee, S., Royce, T., and Tanner, S. (2016). Canvas: versatile and scalable detection of copy number variants. Bioinformatics 32, 2375–2377.

24. Chen, X., Schulz-Trieglaff, O., Shaw, R., Barnes, B., Schlesinger, F., Kallberg, M., Cox, A.J., Kruglyak, S., and Saunders, C.T. (2016). Manta: rapid detection of structural variants and indels for germline and cancer sequencing applications. Bioinformatics 32, 1220–1222.

25. Zarrei, M., MacDonald, J.R., Merico, D., and Scherer, S.W. (2015). A copy number variation map of the human genome. Nat Rev Genet 16, 172–183.

26. (!!! INVALID CITATION !!! 21).

27. Zhu, T., and Niu, D.K. (2013). Frequency of intron loss correlates with processed pseudogene abundance: a novel strategy to test the reverse transcriptase model of intron loss. BMC Biol 11, 23.

28. Thorvaldsdottir, H., Robinson, J.T., and Mesirov, J.P. (2013). Integrative Genomics Viewer (IGV): high-performance genomics data visualization and exploration. Brief Bioinform 14, 178–192.

29. Zhao, X., Emery, S.B., Myers, B., Kidd, J.M., and Mills, R.E. (2016). Resolving complex structural genomic rearrangements using a randomized approach. Genome Biol 17, 126.

30. Wieczorek, D., Bogershausen, N., Beleggia, F., Steiner-Haldenstatt, S., Pohl, E., Li, Y., Milz, E., Martin, M., Thiele, H., Altmuller, J., et al. (2013). A comprehensive molecular study on Coffin-Siris and Nicolaides-Baraitser syndromes identifies a broad molecular and clinical spectrum converging on altered chromatin remodeling. Hum Mol Genet 22, 5121–5135.

31. Lek, M., Karczewski, K.J., Minikel, E.V., Samocha, K.E., Banks, E., Fennell, T., O’Donnell-Luria, A.H., Ware, J.S., Hill, A.J., Cummings, B.B., et al. (2016). Analysis of protein-coding genetic variation in 60,706 humans. Nature 536, 285–291.

32. Richards, S., Aziz, N., Bale, S., Bick, D., Das, S., Gastier-Foster, J., Grody, W.W., Hegde, M., Lyon, E., Spector, E., et al. (2015). Standards and guidelines for the interpretation of sequence variants: a joint consensus recommendation of the American College of Medical Genetics and Genomics and the Association for Molecular Pathology. Genet Med 17, 405–424.

33. Bramswig, N.C., Ludecke, H.J., Hamdan, F.F., Altmuller, J., Beleggia, F., Elcioglu, N.H., Freyer, C., Gerkes, E.H., Demirkol, Y.K., Knupp, K.G., et al. (2017). Heterozygous HNRNPU variants cause early onset epilepsy and severe intellectual disability. Hum Genet 136, 821–834.

34. Namburi, P., Ratnapriya, R., Khateb, S., Lazar, C.H., Kinarty, Y., Obolensky, A., Erdinest, I., Marks-Ohana, D., Pras, E., Ben-Yosef, T., et al. (2016). Bi-allelic Truncating Mutations in CEP78, Encoding Centrosomal Protein 78, Cause Cone–Rod Degeneration with Sensorineural Hearing Loss. Am J Hum Genet 99, 777–784.

35. Nikopoulos, K., Farinelli, P., Giangreco, B., Tsika, C., Royer-Bertrand, B., Mbefo, M.K., Bedoni, N., Kjellstrom, U., El Zaoui, I., Di Gioia, S.A., et al. (2016). Mutations in CEP78 Cause Cone-Rod Dystrophy and Hearing Loss Associated with Primary-Cilia Defects. Am J Hum Genet 99, 770–776.

36. Sedlazeck, F.J., Rescheneder, P., Smolka, M., Fang, H., Nattestad, M., von Haeseler, A., and Schatz, M. (2017). Accurate detection of complex structural variations using single molecule sequencing. bioRxiv.

37. Kielbasa, S.M., Wan, R., Sato, K., Horton, P., and Frith, M.C. (2011). Adaptive seeds tame genomic sequence comparison. Genome Res 21, 487–493.

38. Cretu Stancu, M., van Roosmalen, M.J., Renkens, I., Nieboer, M.M., Middelkamp, S., de Ligt, J., Pregno, G., Giachino, D., Mandrile, G., Espejo Valle-Inclan, J., et al. (2017). Mapping and phasing of structural variation in patient genomes using nanopore sequencing. Nat Commun 8, 1326.

39. Liu, P., Erez, A., Nagamani, S.C., Dhar, S.U., Kolodziejska, K.E., Dharmadhikari, A.V., Cooper, M.L., Wiszniewska, J., Zhang, F., Withers, M.A., et al. (2011). Chromosome catastrophes involve replication mechanisms generating complex genomic rearrangements. Cell 146, 889–903.

40. Lee, J.A., Carvalho, C.M., and Lupski, J.R. (2007). A DNA replication mechanism for generating nonrecurrent rearrangements associated with genomic disorders. Cell 131, 1235–1247.

41. Zhang, F., Khajavi, M., Connolly, A.M., Towne, C.F., Batish, S.D., and Lupski, J.R. (2009). The DNA replication FoSTeS/MMBIR mechanism can generate genomic, genic and exonic complex rearrangements in humans. Nat Genet 41, 849–853.

42. Cooper, G.M., Coe, B.P., Girirajan, S., Rosenfeld, J.A., Vu, T.H., Baker, C., Williams, C., Stalker, H., Hamid, R., Hannig, V., et al. (2011). A copy number variation morbidity map of developmental delay. Nat Genet 43, 838–846.

43. Miller, D.T., Adam, M.P., Aradhya, S., Biesecker, L.G., Brothman, A.R., Carter, N.P., Church, D.M., Crolla, J.A., Eichler, E.E., Epstein, C.J., et al. (2010). Consensus statement: chromosomal microarray is a first-tier clinical diagnostic test for individuals with developmental disabilities or congenital anomalies. Am J Hum Genet 86, 749–764.

44. Merker, J.D., Wenger, A.M., Sneddon, T., Grove, M., Zappala, Z., Fresard, L., Waggott, D., Utiramerur, S., Hou, Y., Smith, K.S., et al. (2018). Long-read genome sequencing identifies causal structural variation in a Mendelian disease. Genet Med 20, 159–163.

45. Huddleston, J., Chaisson, M.J.P., Steinberg, K.M., Warren, W., Hoekzema, K., Gordon, D., Graves-Lindsay, T.A., Munson, K.M., Kronenberg, Z.N., Vives, L., et al. (2017). Discovery and genotyping of structural variation from long-read haploid genome sequence data. Genome Res 27, 677–685.

46. Gong, L., Wong, C.-H., Cheng, W.-C., Tjong, H., Menghi, F., Ngan, C.Y., Liu, E.T., and Wei, C.-L. (2017). Nanopore Sequencing Reveals High-Resolution Structural Variation in the Cancer Genome. bioRxiv.

47. Carvalho, C.M., Pehlivan, D., Ramocki, M.B., Fang, P., Alleva, B., Franco, L.M., Belmont, J.W., Hastings, P.J., and Lupski, J.R. (2013). Replicative mechanisms for CNV formation are error prone. Nat Genet 45, 1319–1326.

48. Kong, A., Frigge, M.L., Masson, G., Besenbacher, S., Sulem, P., Magnusson, G., Gudjonsson, S.A., Sigurdsson, A., Jonasdottir, A., Jonasdottir, A., et al. (2012). Rate of de novo mutations and the importance of father’s age to disease risk. Nature 488, 471–475.

49. Goldmann, J.M., Wong, W.S., Pinelli, M., Farrah, T., Bodian, D., Stittrich, A.B., Glusman, G., Vissers, L.E., Hoischen, A., Roach, J.C., et al. (2016). Parent-of-origin-specific signatures of de novo mutations. Nat Genet 48, 935–939.

50. Flottmann, R., Kragesteen, B.K., Geuer, S., Socha, M., Allou, L., Sowinska-Seidler, A., Bosquillon de Jarcy, L., Wagner, J., Jamsheer, A., Oehl-Jaschkowitz, B., et al. (2017). Noncoding copy-number variations are associated with congenital limb malformation. Genet Med.

51. Collins, R.L., Stone, M.R., Brand, H., Glessner, J.T., and Talkowski, M.E. (2016). CNView: a visualization and annotation tool for copy number variation from whole-genome sequencing. bioRxiv.

52. Ordulu, Z., Wong, K.E., Currall, B.B., Ivanov, A.R., Pereira, S., Althari, S., Gusella, J.F., Talkowski, M.E., and Morton, C.C. (2014). Describing sequencing results of structural chromosome rearrangements with a suggested next-generation cytogenetic nomenclature. Am J Hum Genet 94, 695–709.

